# Temperature-dependent and sex-specific phenotypic plasticity in *Heliconius charithonia*

**DOI:** 10.1101/2024.10.22.619625

**Authors:** Fernando Seixas, Sammy Guerrero, Tianzhu Xiong, James Mallet

## Abstract

External environmental cues can influence the development trajectory of organisms and lead to the formation of distinct phenotypes from the same genotype. Phenotypic plasticity is common in Lepidoptera, with temperature frequently playing a key role. The zebra-longwing (*Heliconius charithonia*) gets its name from its characteristic black wings with yellow stripes. In some populations, yellow scales are sometimes partially replaced by orange (‘rusty’) scales. The proportion of individuals with rusty scales varies geographically and individuals within the same population can vary drastically for this phenotype. Whether such variation is genetically encoded, or environment-dependent is not known. Here we test the influence of temperature in producing the rusty scales phenotype, as well as on the developmental rates, weight, and mortality in this species. Only female butterflies reared in warm controlled conditions develop rusty scales on the yellow parts of their wings. However, under more extreme (warmer) conditions, both males and females exhibit pronounced rusty scale phenotypes, indicating a sexually dimorphic response to temperature that affects both sexes. This temperature-sensitive trait can be triggered during both larval and pupal stages. Temperature also affects life-history traits, and we observed a trade-off between developmental rate and mortality. Individuals raised in warmer conditions develop more rapidly than those in cooler environments, but this accelerated growth is accompanied by higher mortality across all developmental stages.

## Introduction

Phenotypic plasticity is the property whereby a single genotype produces distinct phenotypes depending on environmental conditions (Stearns, 1989) and can be a major driver of phenotypic diversity (Whitman & Agrawal, 2009). Environmental conditions can have direct or trans-generational effects on an individual’s phenotype. Within the same generation, changes in adult phenotypes can result from changes in the environment of the adult (acclimation, often reversible in adult phenotypes), or depend on the conditions experienced during development (developmental plasticity, often irreversible in adult phenotypes).

In moths and butterflies (Lepidoptera), larval and early pupation stages are critical periods for determining wing patterns (Brakefield, 1996; Brakefield et al., 1996). Environmental conditions experienced during these periods (i.e. developmental plasticity) such as temperature, photoperiod, and humidity play a significant role in plasticity of wing color patterns in butterflies (Brakefield, 1996; Kooi & Brakefield, 1999). For instance, increased melanization of wing patterns results from rearing at lower temperatures (Otaki, 2008; Otaki et al., 2010; Sourakov, 2017).

Phenotypic plasticity in wing patterns is particularly well documented in species with seasonal polyphenism (Brakefield & Larsen, 1984; Shapiro, 1976). In the Satyrine butterfly *Bicyclus anynana*, temperature plays a critical role in seasonal variation of eye spots and wing coloration (Brakefield & Reitsma, 1991). In cooler, wet-season environments, butterflies develop smaller eye spots and darker wing colors, which aid in camouflage and predator avoidance. In contrast, during the warmer, dry season, *B. anynana* exhibits larger eye spots and lighter wing coloration, which are thought to play a role in mate attraction.

The passion-vine butterflies, *Heliconius* (Nymphalidae: Heliconiinae), are well known for striking phenotypic diversity, with drastic wing pattern differences between races and/or closely related species, and Müllerian mimicry convergence to distantly related species (Jiggins, 2016). This diversity in wing patterns can be explained by genetic variation mainly at a few major loci (Martin et al., 2013; Moest et al., 2020; Nadeau et al., 2014). However, there tends to be little phenotypic diversity in wing pattern among individuals of the same geographic form in the wild, hence phenotypic plasticity is thought to have little or no effect on the expression of *Heliconius* wing patterns (Jiggins, 2016).

Within this genus, the zebra-longwing (*Heliconius charithonia*) has a widespread distribution, spanning from South to North America, and the Greater Antilles. This species has characteristic black wings with yellow stripes and is phenotypically homogenous throughout its range (Comstock & Brown, 1950a). In some populations, some individuals have orange (‘rusty’) scales overlying the yellow parts of the wing, a phenotype almost limited to females (Comstock & Brown, 1950a). The proportion of individuals with rusty scales varies geographically, and the prevalence of rusty scales can also vary among individuals within the same population. It is not known whether variation in this phenotype is correlated with genetic variation or is driven by environmental cues during development. In this study we test how variation in temperature affects the ‘rusty’ scales phenotype and explore the effects of temperature on life-history traits at different stages of development.

## Materials and Methods

### Controlled rearing experiment

Captive-bred populations of *H. charithonia tuckeri* from Florida were reared in greenhouses. Adult butterflies were kept in a cage and fed using artificial flowers with sugar solution supplemented with pollen and provided additional pollen sources – *Lantana spp*. (Verbenaceae), but without larval host plants. Adult females were transferred, one at a time, into a new cage together with two sets of shoots of a larval host-plant, *Passiflora biflora*, on which females would lay eggs. To guarantee that each shoot received similar numbers of eggs if a female tended to favor one of the shoots, we temporarily isolated the favored shoot in a small rearing cage, allowing the female to oviposit on the other shoot. Once the female oviposited at least 6 eggs on each shoot, she was marked and returned to the main cage, and the day of oviposition was recorded. The two sets of eggs were then placed in separate growth chambers at different temperatures – cool (22ºC) or warm (30ºC). Other conditions were identical in both growth chambers – humidity (80%) and light period (11 hours of light from 7 AM to 6 PM). Caterpillars were fed regularly with new *Passiflora biflora* shoots. To explore stage-specific responses to temperature, the same experiment was conducted but the conditions were switched on pupation (i.e. caterpillars reared in the cool chamber were moved to the warm chamber within 24 hours of pupation, and *vice versa*).

### Phenotype measurements

Durations of larval and pupal periods were recorded in days. Within 24 hours of emergence, adults were sexed and weighed. Each individual was weighed on a Hochoice ES05B analytical balance, with 0.001g resolution. Butterfly wings were observed under a Leica EZ4HD microscope and/or a TOMLOV Digital Microscope for the presence or absence of rusty scales and photographed with a Canon EOS 600D DSLR camera.

## Results

### Elevated temperature induces rusty scales phenotype

We found that wing coloration was sex-specific and plastic. All females reared in warm conditions showed a high density of rusty scales (4 out of 4) while those reared in cool conditions presented a very reduced number or absence of rusty scales (11 out of 11). There was a significant association between rearing conditions and rusty scales phenotypes in females (Fisher exact test, P=0.0007). In males, rearing conditions had no effect on phenotype, with no individuals showing more than a very small number of rusty scales independent of rearing conditions (5 out of 5 and 4 out of 4, in cool and warm conditions respectively).

In switched conditions, 4 out of 7 (57%) females initially reared in cool conditions (CW), and 2 out of 3 (67%) females initially reared in warm conditions (WC) developed rusty scales. However, the proportion of rusty scales was lower when compared to that of females reared in warm conditions during both larval and pupal stages. In males, rusty scales again did not develop, regardless of the conditions.

These results show that the rusty scales phenotype is temperature-dependent and specific to females. Furthermore, temperature cues during both larval and pupal stages contribute to the phenotype, and rusty scales can still develop even in the absence of warm conditions in one of these stages, although to a lesser degree.

### Developmental time – faster development in warm conditions

Median developmental time across all individuals was 10.5 days shorter in individuals reared strictly under cool conditions (CC: 31 days) compared to those reared under warm conditions (WW: 20.5 days) – Figure 2a; Supplementary Table 1. However, this effect was more pronounced in females, for which a reduction in developmental time was observed during both larval and pupal stages, while in males only the pupal stage was clearly affected by rearing conditions (Figure 2b,c). In females, median egg, larval and pupae developmental time were each 4 days shorter (Figure 2b,c; Supplementary Table 1). In males, median larval developmental time was similar under both conditions, but with higher variance for individuals reared under warm conditions (Figure 2b; Supplementary Table 1). Male median pupal developmental time was 3 days shorter under warm conditions.

**Figure 1.**
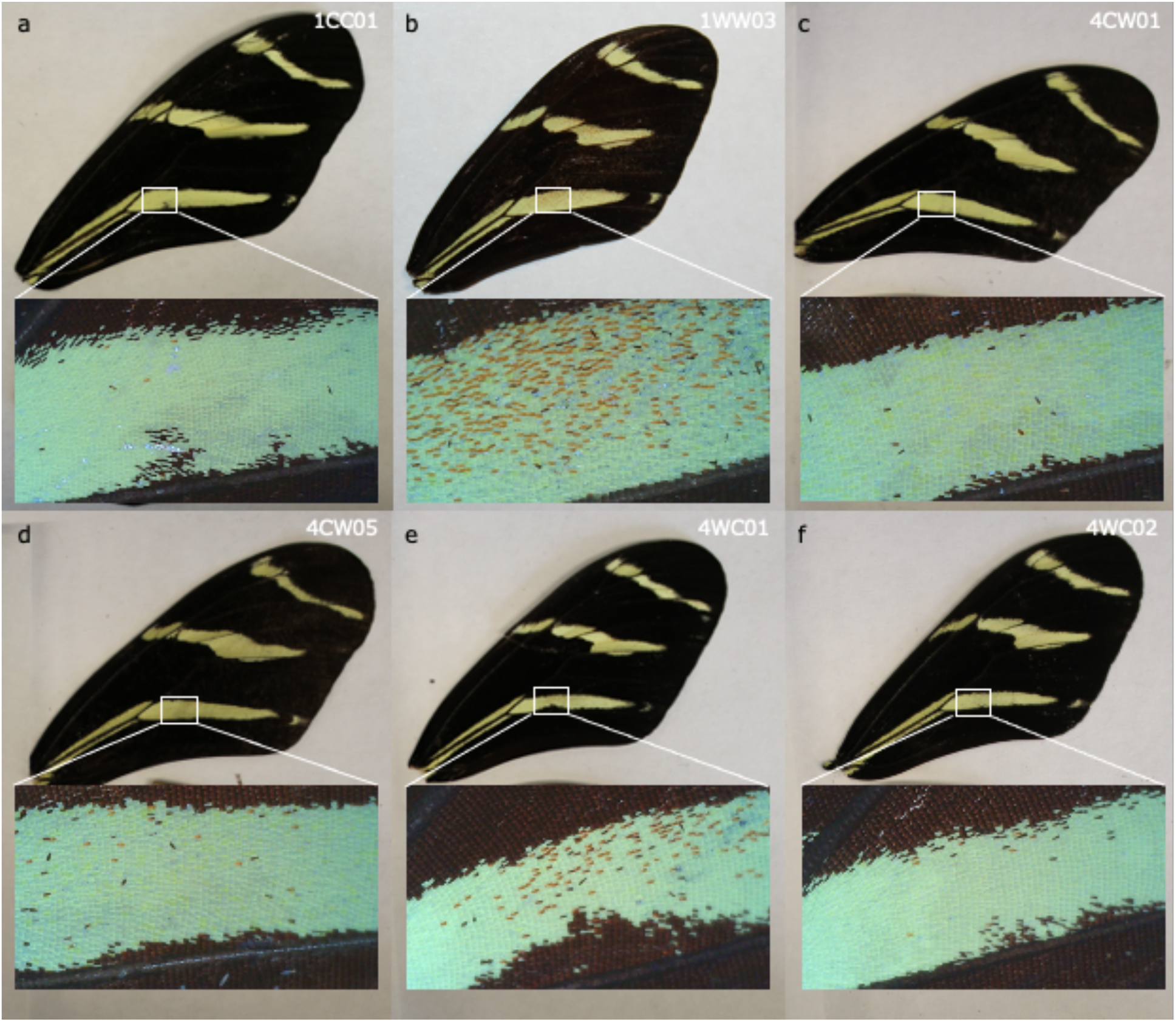
Effect of temperature on the presence/absence of rusty scales in females. Females reared in cool (a) and warm (b) conditions through both larval and pupal stages. Females reared in alternate conditions - cool + warm (c-d) and warm + cool (e-f) - during larval and pupal stages, respectively. Individual codes are given in each panel (see Supplementary Table 1).

**Figure 2.**
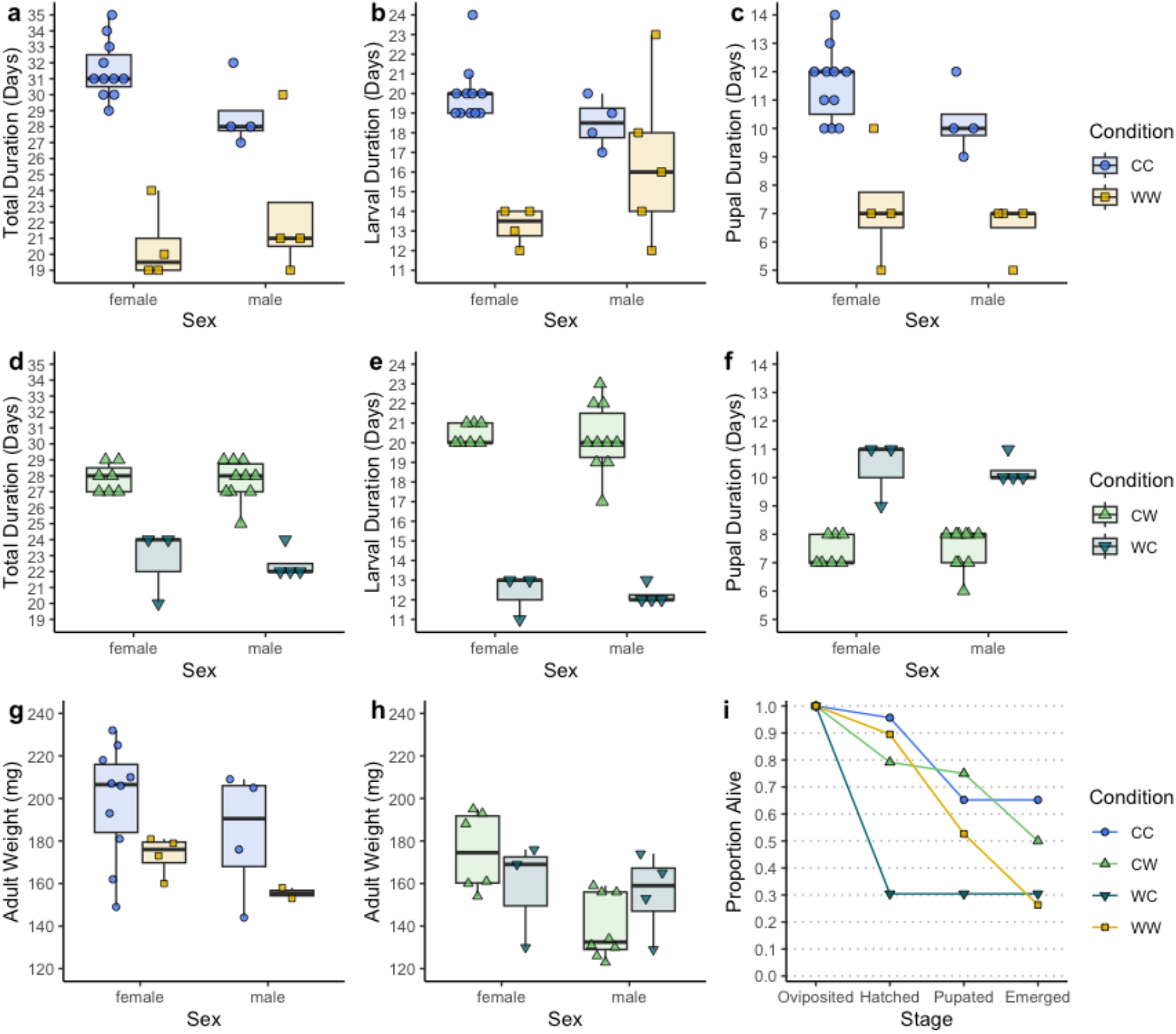
Effect of temperature on life-history traits. Effect of temperature on developmental time (**a-c**) under a single condition throughout both larval and pupal stages and (**d-f**) when reared in switched conditions – cool to warm (CW) and warm to cool (WC). (**g**,**h**) Effect of temperature on adult weight. **i**. Mortality rates throughout the different developmental stages.

Median total developmental time was 4-6 days shorter when larvae reared in warm conditions were then switched to cool (WC), for both sexes (Figure 2d). Larval duration was 5-6.5 days shorter for caterpillars reared under warmer conditions Figure 2e), and pupal duration was also shorter (2-4 days) under warm conditions, independently of how larvae were raised (Figure 2f).

### Adult Weight – no effect of temperature on adult weight

There were no significant differences in adult weight between rearing conditions, either in females (Wilcoxon rank sum test, p-value = 0.1561) or males (Wilcoxon rank sum test, p-value = 0.5333), although there was increased variation when reared under cooler conditions in both sexes (Figure 2g). There were no significant differences in weight between the sexes when reared under the same conditions (Wilcoxon rank sum test, p-value = 0.2529) – Figure 2g.

After switching conditions on pupation, initial conditions had no significant effect on adult weight, either of females or males (Figure 2h). When reared initially in cool conditions and changed to warm conditions upon pupation, females had significantly higher adult weights than males (Wilcoxon rank sum test, p-value = 0.004022), but there was no significant difference if initially reared in warm conditions (Wilcoxon rank sum test, p-value = 0.6286) - Figure 2h.

### Mortality Rates – higher mortality rates in warmer conditions

In treatments with constant temperature, mortality rates were higher under warmer conditions throughout all stages of development (78.3 %) compared to cool conditions (34.8%) – (Figure 2i, Supplementary Table 1). When in cool conditions, 34.8% of individuals died during the egg (4.3%) and caterpillar (30.4%) stages and all pupae eclosed as adults. On the other hand, when reared in warm conditions, deaths occurred during all stages of development.

Switched conditions allow one to disentangle the interplay between developmental stage and temperature in mortality rate (Figure 2i). Overall, the mortality rate (i.e., the proportion of healthy butterflies emerging relative to the number of eggs oviposited) was higher when caterpillars were initially reared in warm conditions (69.6%) compared to when individuals were initially reared in cool conditions (50.0%). When initially reared in warm conditions and switched to cool conditions, all deaths happened at the eggs/caterpillar stage (69.6% of individuals died), all surviving individuals making it to the pupal and adult stage. On the other hand, when individuals were reared initially in cool conditions, only 20.8% of individuals failed to hatch and only 25% failed to reach the pupal stage. Of those reaching the pupal stage, 29.4% failed to emerge or emerged with malformations. These results show that the highest mortality occurs under warmer conditions, regardless of developmental stage.

Comparing life stages under the same conditions, eggs failed to hatch more often under warm conditions (WW: 11.8% and WC: 60.6% vs. CC: 4.3% and CW: 20.8%), but for larvae there was an opposite trend (WW: 17.6% and WC: 0.0% vs. CC: 30.4% and CW: 4.2%) - Figure 2i. Pupae reared under cool conditions never died and adults emerged without malformations, independently of initial conditions (CC: 0%, CW: 0%). Pupae reared under warm conditions often died or eclosed malformed. This was independent of rearing conditions of eggs and larvae (WW: 17.6% and WC: 16.7%). Overall, while deaths are expected during the egg and larval stages regardless of temperature, failure to emerge or malformation happened only under warm conditions (Figure 2i).

Finally, egg density in the small cages also seems to have an effect on mortality rates. Families reared under switched conditions experiments happened to have larger batch sizes (n=9-14) and higher mortality (CW: 20.8% and WC: 69.6%), compared to those reared under stable conditions (n=4-9; CC: 4.3% and WW: 11.8%) – (Supplementary Table 1).

## Discussion

Phenotypic plasticity in *Heliconius* has been documented for several traits, including development time, larval survival, adult size, wing shape and brain morphology (Hebberecht et al., 2023; Huang et al., 2024; Rodrigues & Moreira, 2002, 2004) but, unlike in many other butterflies, wing color phenotype plasticity has not been noted (Jiggins, 2016). In this study, we provide experimental evidence that temperature during various developmental stages, such as egg, larval, and pupal phases, not only significantly influences life-history traits in the zebra-longwing, *H. charithonia*, but also color of wing scales.

Variation in wing coloration and life-history traits in Lepidoptera can be environmentally induced through different environmental cues (Otaki, 2008; Shimajiri & Otaki, 2022; Sourakov, 2017). In this study, we focused on the effects of temperature, but it is likely that other environmental factors (e.g. humidity and day length) could also affect these traits, and possibly lead to more extreme phenotypes particularly regarding wing colors patterns. In the greenhouse, temperature was regulated by a combination of air conditioning and vents which allow inside temperature to match outside temperature. Temperature was generally well maintained within defined parameters (25-28°C), while day length followed the natural conditions, and humidity was controlled by watering the floors. The combination of these factors could thus vary in our stock populations, and perhaps as a result, we observed some individuals with more extreme rusty scale phenotypes than in our controlled experiments (Figure 3a). Individuals with similarly darker orange phenotypes have also been observed in the wild (see Sourakov et al., 2018), probably as a result of variation in environmental conditions.

**Figure 3.**
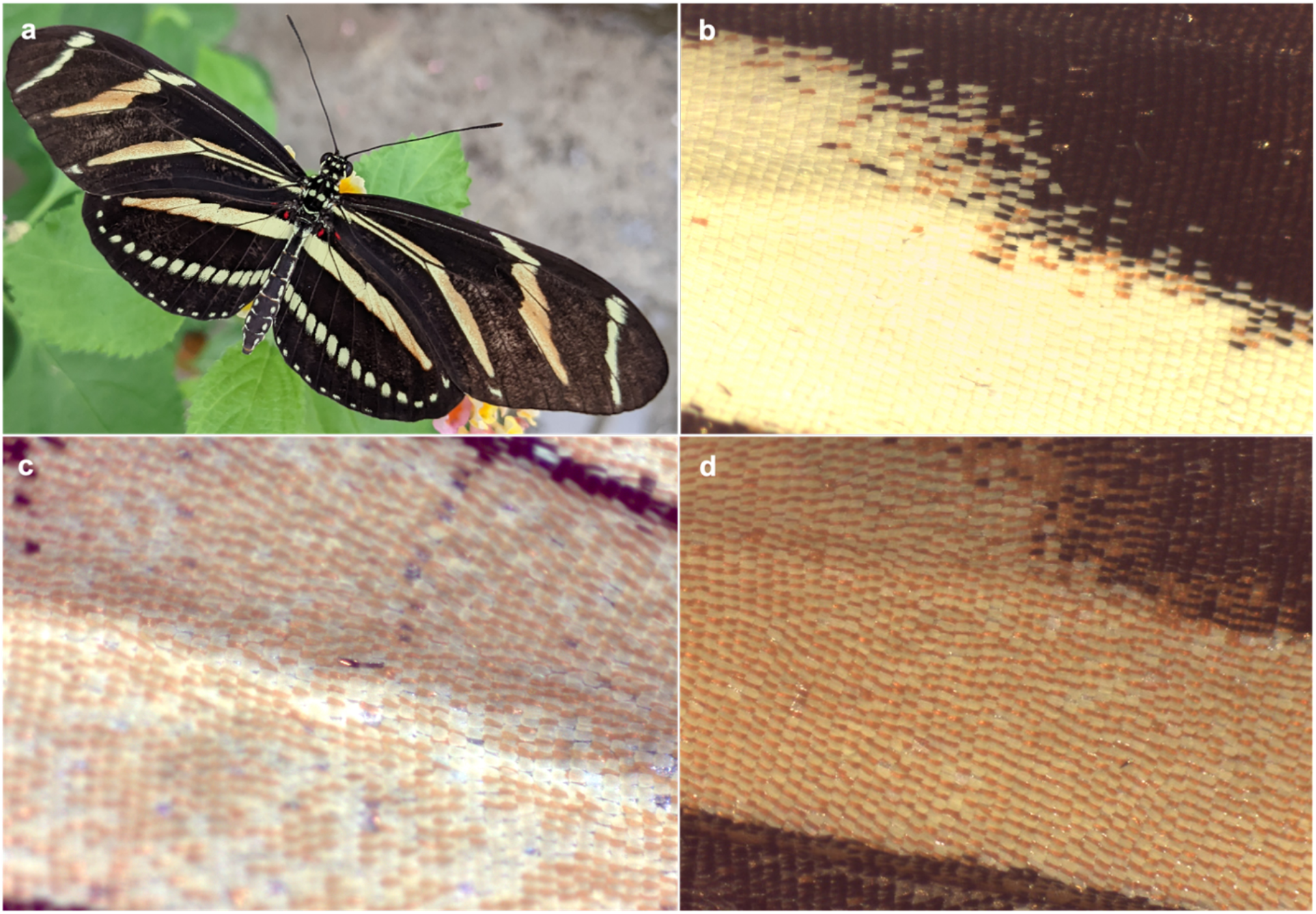
Extreme rusty scale phenotypes in greenhouse stock populations. **a**. *H. charithonia tuckeri* female from (Florida) **b**. *H. charithonia bassleri* male (Colombia) **c**. *H. charithonia vasquezae* female (Texas) Texas **d**. *H. charithonia bassleri* female (Colombia).

However, the most extreme phenotypes in our greenhouse populations were observed as a result of a chance event: failure of the air conditioning system in one of our greenhouses during a brief period in summer). During that period, pupae imported from Colombia (*H. c. bassleri*) and larvae and pupae from our captive bred Texas population (*H. c. vasquezae*) developed under temperatures that occasionally reached 40°C. While many individuals failed to emerge, the yellow wing bands of those that survived were extensively covered by rusty scales (Figure 3b-d). These extreme phenotypes are similar to those observed by Sourakov et al. (2018), in individuals reared in the greenhouse from a batch of eggs collected in the wild although, in that case, the extreme dark phenotypes likely resulted from a baculovirus infection (Sourakov et al., 2018). In our controlled experiments, only females showed rusty scales, but they developed also in males subjected to these extreme conditions in the greenhouse (Figure 3b). In natural populations, the rusty scales phenotype is also found in both sexes but the proportion of females with this phenotype is much larger than in males: 35.2% of females had rusty scales, compared to only 1.2% of males, across all samples and populations (Comstock & Brown, 1950b). The higher prevalence of rusty scales in females in our experiments and in natural populations suggests sex-specific reaction norms to temperature in the production of ‘rusty’ scales.

Plasticity and sexual dimorphism in other wing traits – wing shape and size – has also been observed in natural populations of *H. charithonia* (Ramos-Pérez et al., 2020). Wing morphology, and particularly wing color patterns, are generally under strong selection in *Heliconius* due to their functions in aposematism and mimicry (Jiggins, 2016). Accordingly, since deviations from the optimum mimicry phenotype are selected against, *Heliconius* tend to show no or very limited sexual dimorphism and plasticity in wing phenotypes locally (Jiggins, 2016). Unlike other *Heliconius, H. charithonia* is non-mimetic. Hence selection on these traits is hypothesized to depend only on local density of this species, and this might allow phenotypic plasticity to evolve despite any adaptive significance. Instead, the rusty scale phenotype might be simply a by-product of differential expression of a gene(s) involved in adaptation to extreme temperatures also involved in scale color pathways. For instance, trehalose is a sugar found in the hemolymph of insects and is involved in adaptation to extreme temperatures and desiccation (Andersen et al., 2011; Benoit et al., 2009; Elnitsky et al., 2008). This is achieved either through the upregulation of genes responsible for converting trehalose into glucose or facilitating its transport between cells and the hemolymph (Kikawada et al., 2007; Liu et al., 2013; Zhou et al., 2022). Genes involved in the transport of trehalose have also been suggested to be implicated in adaptation to colder temperatures and desiccation in *H. charithonia* in Northern parts of its range (Seixas et al., 2024). Concurrently, increased expression of the *trehalase* gene, which encodes an enzyme that converts trehalose into glucose, has been linked to the production of red pigmentation in *Junonia coenia* reared under warm conditions (van der Burg et al., 2020). It is therefore plausible that in *H. charithonia*, genes regulating trehalose conversion or transport are differentially expressed in response to elevated temperatures, promoting thermal adaptation while incidentally producing the rusty scale phenotype. However, since this species is aposematic, we cannot discard the possibility that individuals presenting the rusty scales phenotype also incur into a fitness cost from increased predation.

Finally, temperature is also a key factor influencing numerous life history traits in Lepidoptera, including development rate, body size and mortality (Rodrigues & Beldade, 2020). Trade-offs frequently occur between these traits, where a favorable change in one trait is offset by negative effects on another, reflecting a balance of fitness costs (Stearns, 1989). For instance, exposure to high temperatures often accelerates growth but can reduce fitness and lead to mortality, primarily through protein denaturation, membrane destabilization, and desiccation (Chown & Terblanche, 2006; Klose & Robertson, 2004; Potter et al., 2009). Under elevated temperatures, there may also be a trade-off between developmental time and morphology, resulting in emergence at a lower adult body size (Chown & Terblanche, 2006). In line with previous studies, including in *Heliconius* (Huang et al., 2024), we observe trade-off effects of temperature in developmental rates and survival rates. Individuals reared at higher temperature had faster developmental rates but at the cost of lower survival, at all stages of development. However, our results indicate that temperature does not affect adult weight. Similarly, a study on *Heliconius erato phyllis* (Rodrigues & Moreira, 2004) found that temperature during larval development had no impact on adult size. Instead, adult size was primarily determined by the type, availability, and quality of host plants consumed during the larval stage (Rodrigues & Moreira, 2002). This suggests that temperature is likely not a critical factor in determining size in *Heliconius* species.

## Author’s contributions

FS designed this study. FS, JM and SM tended to the plants and reared butterflies, while FS and SM carried out experimental work, and measurements, with the assistance of TX. FS and JM wrote the manuscript.

## Acknowledgments

We thank Elena Mistreanu for assisting in rearing of butterflies. We thank Neil Rosser for collecting individuals that we used to establish a stock population of Florida *H. charithonia*.

